# Mitochondrial DNA variants segregate during human preimplantation development into genetically different cell lineages that are maintained postnatally

**DOI:** 10.1101/2021.11.05.467445

**Authors:** Joke Mertens, Marius Regin, Neelke De Munck, Edouard Couvreu de Deckersberg, Florence Belva, Karen Sermon, Herman Tournaye, Christophe Blockeel, Hilde Van de Velde, Claudia Spits

## Abstract

Humans present remarkable mitochondrial DNA (mtDNA) variant mosaicism, not only across tissues but even across individual cells within one person. The timing of the first appearance of this mosaicism has not yet been established. In this study, we hypothesized it occurs during preimplantation development. To investigate this, we deep-sequenced the mtDNA of 254 oocytes from 85 donors, 158 single blastomeres of 25 day-3 embryos, 17 inner cell mass and trophectoderm samples of 7 day-5 blastocysts, 142 bulk DNA and 68 single cells of different adult tissues. We found that day-3 preimplantation embryos already present blastomeres that carry variants unique to that cell, showing that the first events of mtDNA mosaicism happen very early in human development. We classified the mtDNA variants based on their recurrence or uniqueness across sibling oocytes and embryos, and between single cells and samples from the same embryos or adult individuals. Variants that recurred across samples had higher heteroplasmic loads and more frequently resulted in synonymous changes or were located in non-coding regions than variants that were unique to one oocyte or single embryonic cell. These differences were maintained through developmental stages, suggesting that the mtDNA mosaicism arising in preimplantation development is maintained into adulthood. Further, the results support a model in which close clustering of mitochondria carrying specific mtDNA variants in the ooplasm leads to asymmetric distribution of these mitochondria throughout the cell divisions of the preimplantation embryo, resulting in the appearance of the first form of mtDNA mosaicism in human development.

## INTRODUCTION

The vast majority of studies in human genetics have been performed on bulk DNA, extracted from peripheral blood or other tissues. In recent years it has become increasingly obvious that the cells of our body are not as genetically homogeneous as previously thought. Next to the already classically well-known cases of cellular mosaicism, such as the variation in the somatic rearrangements of immunoglobulin and T-cell receptor in lymphocytes, a whole new dimension of diversity is just being uncovered owing to the emergence of single cell comprehensive genome analysis^1^. The mutations driving somatic mosaicism probably occur at all stages of development, from early preimplantation development, as seen for chromosomal abnormalities^2^, to the ageing individual^1^.

The mitochondrial DNA (mtDNA) is, in this sense, particularly diverse. It is known for long that a given inherited mtDNA mutation can be present in different loads in different individuals of the same family, and in different tissues of the same individual^3^, and somatic cellular heterogeneity has been shown in blood cells^4,5^, neurons, glia^6^, and single muscle fibers^7^. In recent years, the advent of massive parallel sequencing has had a deep impact on the field of mitochondrial genetics. The fact that the relatively small mtDNA can be sequenced at high depth has facilitated the simultaneous detection of all variants within this genome and their individual heteroplasmic loads^8,9^. This type of deep sequencing work has shown, for instance, that pathogenic mtDNA variants are commonly present in healthy individuals, with a mean load of 2% and cross-generation studies have shown that these variants are heritable^8–10^.

The composition and heteroplasmic load of the variants in the mtDNA of an individual can vary during development, as it goes through various bottlenecks. The first bottleneck occurs during oogenesis, where the few mtDNA copies in the primordial germ cells replicate rapidly, having increased a thousand-fold when reaching the mature oocyte stage. During this process, low-load heteroplasmic variants in early primordial germ cells can increase to much higher loads in the late primordial germ cells, where selection mechanisms will eliminate mitochondria with variants affecting their functionality^11^. This selection process circumvents Muller’s ratchet, an evolutionary process where deleterious variants can accumulate rapidly over generations in an irreversible manner due to uniparental inheritance and lack of genomic recombination of the mtDNA^12^. This first bottleneck is also responsible for the diversity in heteroplasmic loads of the same variant across children of the same mother. The second bottleneck occurs after fertilization, where the mtDNA copy number per cell declines transiently during early embryonic development due to halted replication, which is resumed when the embryonic cells initiate differentiation^13,14^. Finally, later in development, the mtDNA becomes susceptible to somatic mutagenesis due to errors of the polymerase gamma, its proximity to reactive oxygen species and a very low protection against mutagenesis by repair mechanisms and histones^15–17^, leading to ageing-related somatic mosaicism (reviewed by van den Ameele et al. in 2020^18^).

Several studies have demonstrated mtDNA mosaicism for both disease and non-disease-causing variants, both at the single-cell level and across tissues of one individual^19–22^. While some of this variation has been attributed to ageing^20^, the recurrence of other mosaic variants across multiple cells of the same individual suggests that the variant emerged very early in development, after or during the formation of the three germ layers. Other variants appear to be tissue-specific, leading to the suggestion that mtDNA variant composition can be cell-type specific^21,22^. Overall, these mosaic mtDNA variants can be present in one cell at a very high load, while in other cells it is at a very low load or not present at all. These differences across cells of the same individual could be explained through clonal expansion by random genetic drift. Because this process is considered to be relatively slow, the initial event has been proposed to occur early in development, but the exact timing and mechanism remain to be elucidated^23^.

Currently, all the knowledge on the segregation of mtDNA variants during early human development is based on the study of inherited pathogenic mtDNA mutations, mostly in the context of preimplantation genetic testing. A significant number of studies have explored the possibility of quantifying pathogenic heteroplasmic variants in polar bodies, single blastomeres biopsied at the cleavage stage or trophectoderm biopsies at the blastocyst stage and have investigated if the results of these biopsies are representative of the rest of the embryo. Work has been published on the segregation of mutations causing mitochondrial diseases such as Leigh syndrome^24,25^, NARP^26^, Lieber’s disease^27^ and MERF/MELAS^28–31^. With some exceptions^31^, the different groups have found a good consistency in heteroplasmic loads between samples of the same embryo, showing that these variants are homogeneously distributed across the mitochondria in the oocyte, and homogeneously partitioned during the early developmental cell divisions. Conversely, there is no knowledge on the appearance and segregation of non-disease causing mtDNA variants during early human development.

In this study, we aimed at determining the timing of appearance mosaicism for non-disease associated mtDNA variants during human preimplantation development. We studied to which extent individual embryonic cells differed from each other and identified different types of mtDNA variants depending on their recurrence across samples of the same donor. Comparison of the patterns of mtDNA variants in oocytes, day-3 and day-5 embryos, and adult-stage tissues and single cells revealed that mtDNA mosaicism appears as early as day 3 of human development, and is maintained through development, resulting in genetically diverse cell lineages in the adult.

## MATERIAL AND METHODS

### Sample and single-cell collection

All buccal (N=59), blood (N=57) and urine (N=26) samples and oocytes (N=254) and embryos on day 3 (n=25) or at the blastocyst stage (n=7) were obtained after signed informed consent from the donors at the Center for Medical Genetics and the Brussels IVF Center for Reproductive Medicine of the UZ Brussel (Supplementary table S1). Supernumerary oocytes were donated after oocyte pick-up while preimplantation embryos were donated for research after the legally determined cryostorage period of 5 years passed^32^. Prior to the start of the study, approval was acquired from the Local Ethical Committee of the Vrije Universiteit Brussel and the Universitair Ziekenhuis Brussel, and by the Federal Ethical Committee on Medical and Scientific Research on embryos in vitro.

Day-3 embryos were warmed using the Vitrification Thaw kit (Vit Kit-Thaw, Irvine Scientific, USA) according to manufacturer’s instructions. Subsequently, they were left to recover in 25 µL droplets of Origio blastocyst medium (Origio, The Netherlands) for 3h in an incubator at 37°C with 89% N_2_, 6% CO_2_ and 5% O_2_. The day-3 embryos and oocytes were freed from their zona pellucida by incubating them in a droplet of pronase (100 mg/100µL human tubal fluid) and by gently pipetting them up and down. The inner cell mass (ICM) and trophectoderm (TE) samples were biopsied from day-5 blastocysts that were diagnosed as affected by a monogenic disease after preimplantation genetic testing, as previously described^19^. The oocytes, embryos, ICM and TE samples and adult single cells were washed three times in Ca^2+^- and Mg^2+^-free medium. The individual blastomeres obtained from cleavage stage embryos dissociated in the Ca^2+^- and Mg^2+^-free medium were washed three additional times before collecting in 2.5 µL ALB (alkaline lysis buffer) as described before^33^. The samples were kept at −20°C until further processing. The bulk DNA of the somatic tissues was extracted using a kit according to manufacturer’s instructions (DNeasy Blood and Tissue, Qiagen).

### mtDNA enrichment and massive parallel sequencing

Before PCR, the single oocytes and blastomeres, ICM/TE biopsies from the day-5 blastocysts and adult cells were incubated at 65°C for 10 min to ensure full lysis of the cells. Long-range PCR^34^ was performed using a primer set to generate amplicons of 13 Kbp (5042f – 1424r). The primer sequences for the primer set were 5’-AGCAGTTCTACCGTACAACC-3’ (forward) and 5’-ATCCACCTTCGACCCTTAAG-3’ (reverse). The amplification was done in a total volume of 50 µL per sample containing 10 µL of LongAmp buffer, 2 µL of Taq DNA polymerase (LongAmp Taq DNA Polymerase kit, New England Biolabs), 7.5 µL of dNTPs (dNTP set, Illustra™), 2 µL of each primer (10 µM), 2.5 µL Tricine (200mM, Sigma-Aldrich) and 21.5 µL H_2_O. The PCR protocol started with an initiation step of 30 sec at 94°C followed by a touchdown of 8 cycles of 15 sec at 94°C, 30 sec at 64°C (−0.4°C per cycle) and 11 min at 61°C, 29-37 cycles were added of 15 sec at 94°C, 30 sec at 61°C and 11 min at 65°C and completed with a final elongation step of 11 min at 65°C. Successful amplification was confirmed using agarose gel electrophoresis (1.5%). After PCR purification with AMPure beads (Beckmann Coulter), library preparation as described in Mertens et al.^34^, was performed using the TruSeq DNA PCR-free Library Preparation kit (Illumina). The amplicons were sheared using a CovarisT^M^ M220 sonicator (Life Technologies), following instrument specification to generate fragments of ± 100 base pairs. The detection of the nucleotide sequence was done on the Illumina NovaSeq6000 platform using the according kit (Illumina).

### Data analysis and bioinformatics processing

The generated data was aligned to the reference genome (NC_012920.1) with BWA-MEM and uploaded to mtDNA server^35^ (v1.1.3) which detected the homoplasmic (>98.5% frequency) and heteroplasmic (<98.5% frequency) single nucleotide variants (SNV) as well as the haplogroup and possible contaminations. Insertions and deletions were detected and SNVs were confirmed by Mutect2^36^ (GATK v3.6 Mutect2). The annotations of the variants were done using MitoWheel and the possible amino-acid changes were identified using MutPred2^37^. A more detailed protocol of the bioinformatic processing and the validation of the full sequencing setup can be found in our previously published work^19,34,38^.

### Statistics

Statistics were performed using the two-tailed Fisher’s exact or the Chi-square test, p-values <0.05 were considered significant.

## RESULTS

### Mitochondrial DNA mosaicism occurs as early as day 3 of human development

We deep-sequenced the mtDNA of 254 oocytes from 85 donors, 158 single blastomeres from 25 day-3 embryos obtained from 9 donors, 17 samples from 7 day-5 blastocysts (7 ICM and 10 TE), 142 adult DNA samples from bulk tissue (59 buccal, 57 blood and 26 urine samples) and 68 single cells from buccal swab (N=37) and urine (N=31) of 3 donors. For the oocytes, we collected at least two samples per donor (either two or more oocytes of the same donor or oocytes and somatic tissues) and for the adult DNA samples we also collected at least two tissues per donor.

We identified heteroplasmic variants in 58.7% of the oocytes, 86.1% of blastomeres of day-3 embryos, in all of the ICM/TE samples of day-5 blastocysts, 47.9% of the adult bulk tissues and 95.6% of adult single cells (Figure 1a). The average heteroplasmic variant load was under 10% for the oocytes and embryos (oocytes: 7.6 ± 13.8%, day-3 blastomeres: 6.1 ± 11.2% and day-5 ICM/TE biopsies: 5.1 ± 5.5%), 20.1 ± 27.6% for the adult bulk tissues and 12.1 ± 13.6% in adult single cells (Figure 1b).

**Figure 1.**
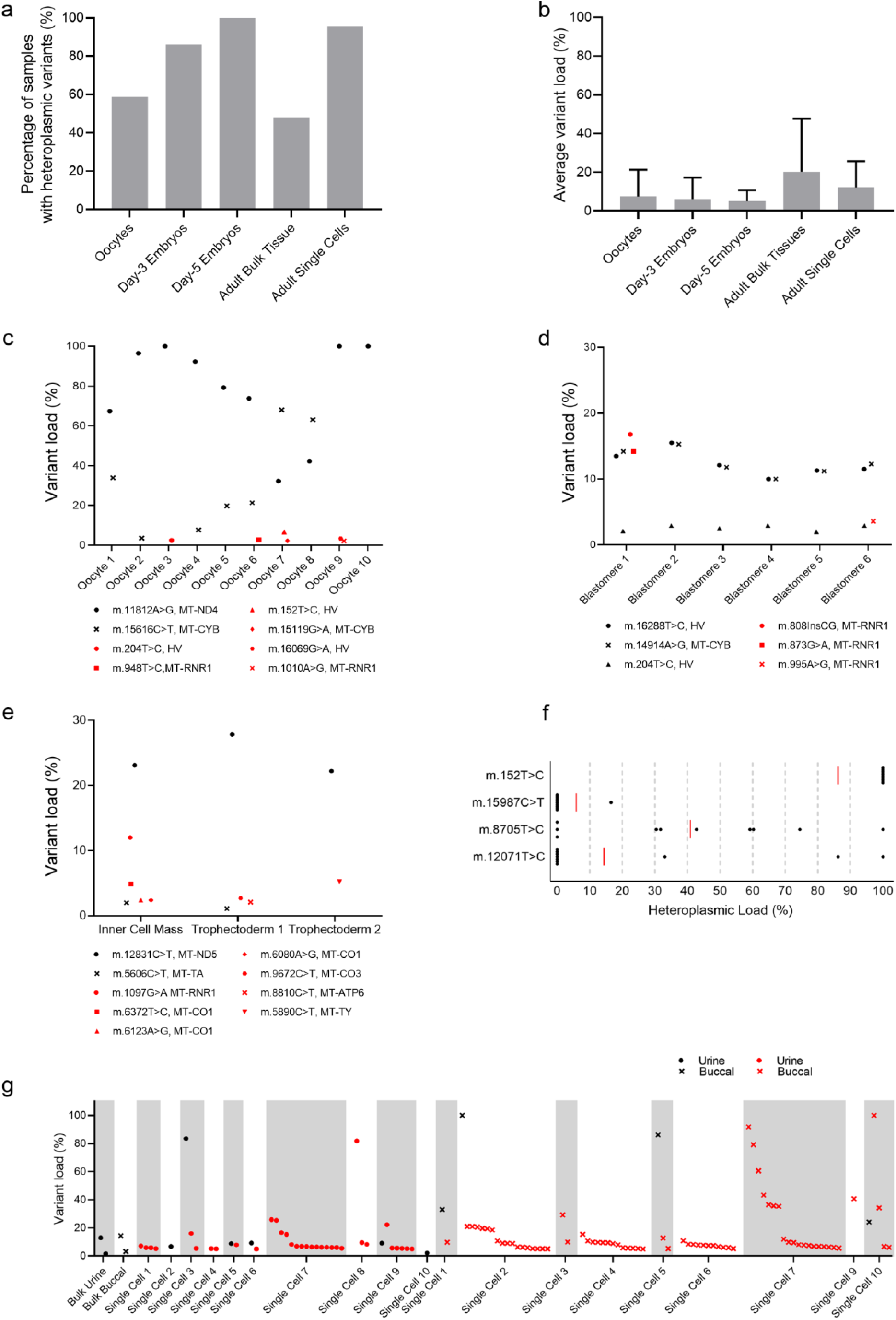
Overview of the heteroplasmic variants found in the different cohorts. **a**. Percentage of samples with heteroplasmic variants in oocytes, day-3 embryos and day-5 blastocysts, adult bulk tissues and adult single cells. **b**. The average load and standard deviation of the heteroplasmic variants found in the oocytes, day-3 embryos and day-5 blastocysts, adult bulk tissues and adult single cells. **c-e:** Examples of variants and their respective load found in oocytes from the same donor (c), blastomeres from the same day-3 embryo (d) and biopsies from the same day-5 blastocysts (e). Variants that recur across cells or samples are shown in black and in red are variants that are unique to one sample. **f**. Heteroplasmic load of recurrent variants found in the bulk DNA sample (red line) and in the single cells from the same tissue (black dots). **g**. Variants found in the adult bulk DNA and in 10 single cells of urine and buccal samples that were recurrent across bulk and single cells (black dots for the urine samples and black crosses for the buccal samples) and variants that were unique to one cell (red dots for the urine cells and crosses for the buccal cells).

Further analysis of the variants showed that while some of the variants recurred across samples of the same donor, other variants were unique to one sample. Therefore, in the further downstream analysis of our dataset, we categorized the heteroplasmic variants as “recurrent” when they were present in multiple tissues of one individual, multiple cells of one day-3 embryo, different biopsies of a day-5 blastocyst or in multiple oocytes of one donor, and as “unique” when they were present in only one sample. An example of the variant composition of different sample types is shown in figure 1c-g. The full datasets can be found in Table S2-S5. We found recurrent mtDNA variants in 26.8% of oocytes, 76.0% of day-3 embryos, 85.7% of day-5 blastocysts, 38.7% of adult bulk DNA samples and 47.1% of adult single cells. Unique variants were found in 48.0% of oocytes, 92.0% of day 3 embryos, 85.7% of day 5 blastocysts, 16.9% of bulk DNA samples and 85.3% single cells.

Overall, the data revealed considerable mtDNA variation across oocytes of the same donor, showing that germ line mosaicism for non-disease associated mtDNA variants is exceedingly common. In the embryos, we found that somatic mtDNA mosaicism occurs already on day-3 of development and is maintained in day-5 blastocysts. This first type of mosaicism is due to the appearance of unique variants in individual cells of the embryo. At this stage of development variants that recur across cells of the embryo do so consistently at very similar loads. These recurrent variants are the source of a second type of somatic mosaicism that emerges later in development and that is evidenced by the results of the adult tissues and single cells. Here, we found that the heteroplasmic load of variants measured in a DNA sample extracted from a given tissue represent the average of the widely variable loads found in the individual cells in that tissue, ranging from a homoplasmic state to absent in some cells (Figure 1f). Finally, a third type of mosaicism is present in the adult single cells, in the form of numerous unique variants that have most likely originated by somatic mutagenesis related to ageing. Interestingly, we have identified four such variants that appear in a tissue-specific manner in different donors: m.215A>G and m.152T>C were found in buccal samples of 7 donors, m.16311T>C in blood and buccal samples of 2 donors and m.72T>C in urine samples of 2 donors.

### Sibling oocytes carry both recurrent and unique variants that differ in location, type and heteroplasmic load

In 26.8% of the oocytes, we found variants that recurred across two or more of the oocytes of the same donor. These variants were evenly distributed between the non-coding and protein-coding regions, and none were detected in the rRNA/tRNA-coding sequences. A non-synonymous change was induced by 16.7% of the variants (Figure 2a). We calculated the mutation rate per base as the number of variants found in each mtDNA locus, divided by the number of sequenced base pairs of that locus. In the oocytes, the highest mutation rate per base was in the hypervariable region (Figure 2b). Of recurrent variants in the oocyte, 61.1% had heteroplasmic loads over 20% (Figure 2c), while their load was independent from their location or type of change (Figure 2d).

**Figure 2.**
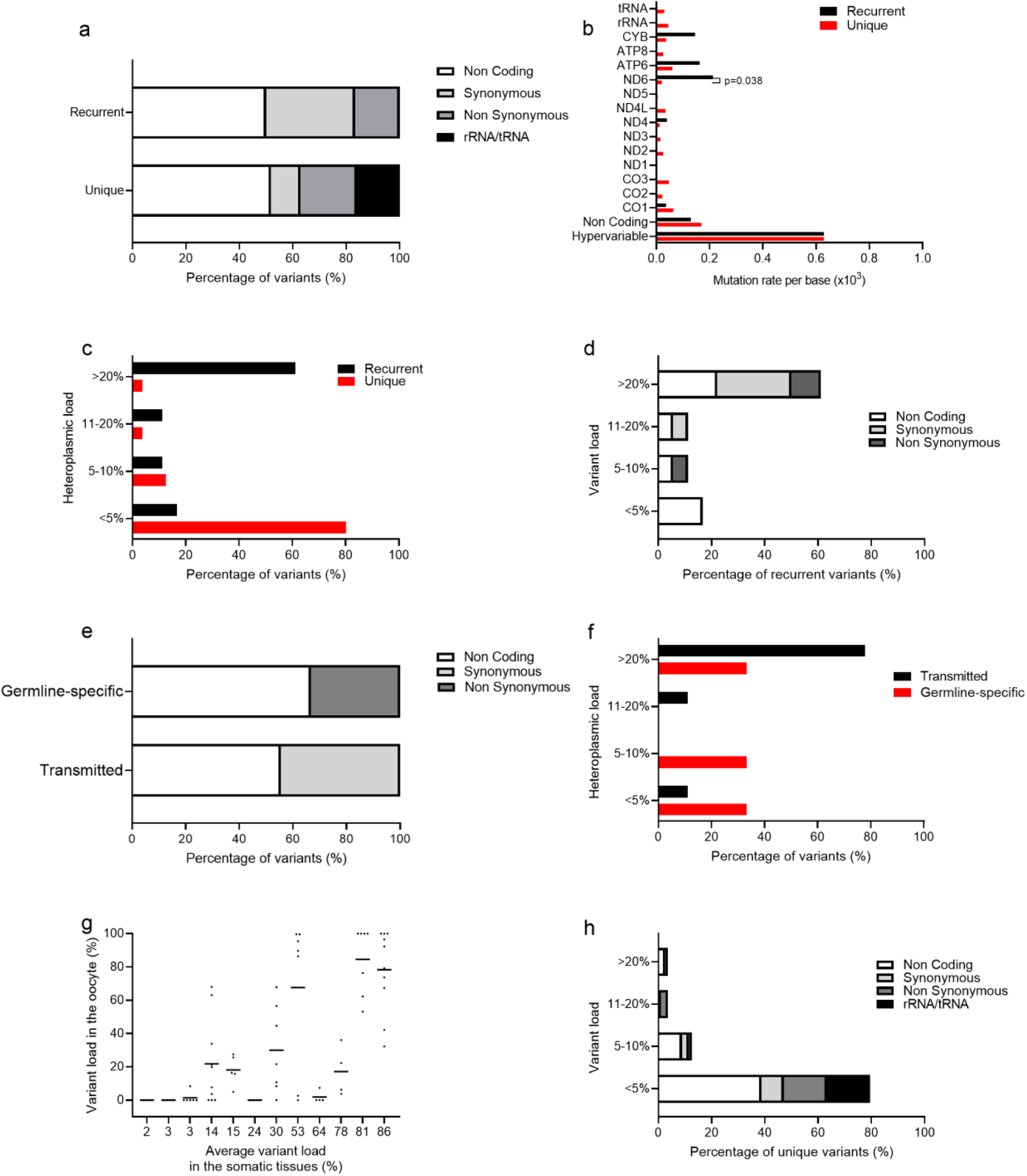
Sibling oocytes carry both recurrent and unique variants that differ in location, type and heteroplasmic load. **a**. Distribution of recurrent and unique variants in oocytes based on their location or type (in case of a variant in the protein-coding regions) in the mitochondrial genome (non-coding, protein-coding synonymous, protein-coding non-synonymous or rRNA/tRNA-coding regions). **b**. Mutation rate per base for the recurrent and unique variants in the oocytes. Variants in the gene *MT-ND6* were more likely to be recurrent (Fisher’s exact test, p=0.038). **c**. Recurrent and unique variants in the oocytes categorized for their variant load. **d**. Recurrent variants in the oocytes categorized for their load and distribution in the mtDNA. **e**. Distribution in the mtDNA based on their location or type of the recurrent variants in the oocytes categorized whether they were present in the somatic tissues of the donor (“Transmitted”) or only present in the oocytes of the donor and not in the somatic tissues (“Germline-specific”). **f**. Recurrent transmitted and germline-specific variants in oocytes categorized for their variant load. **g**. Example of transmitted variants where the average load in the somatic tissues is plotted against the load of the same variant in the oocytes of the respective donor. **h**. Unique variants in the oocytes categorized for their load and distribution in the mtDNA.

We sequenced somatic tissues (buccal swabs, blood and urine) from a subgroup of 25 oocyte donors. In these, 9 variants were present in at least one somatic tissue and were transmitted to the oocytes (referred to as “transmitted”) and 3 variants were only present in the oocytes and not in the somatic tissues (referred to as “germline-specific”). Both types of variants were similarly distributed across non-coding and protein-coding regions with a distribution of 55.6 versus 44.4% for the transmitted variants and 66.7 versus 33.3% for the germline-specific variants. However, the transmitted protein-coding variants were exclusively synonymous while the germline-specific protein-coding variants were all non-synonymous (Figure 2e). Of the transmitted variants, 77.8% were present at loads >20%, while this was only 33.3% for the germline-specific variants, although this difference was not significant (Fisher’s exact test, p=0.2364, Figure 2f). Lastly, the higher the load in the somatic tissues, the higher the likelihood that the variant would be present in the majority of the oocytes of the donor (correlation R=0.65, p<0.0001), with variants with loads as low as 3% in the somatic tissues being identified as well in the oocytes. Conversely, the heteroplasmic load of the variant could significantly differ from oocyte to oocyte, in extreme cases going from undetectable levels in one oocyte to homoplasmy in another (Figure 2g).

Nearly half (48.0%) of oocytes carried variants that were unique to one oocyte in a cohort, and these were remarkably different in their location and heteroplasmic load from the recurrent variants. Of the unique variants, 16.3% were located in the rRNA/tRNA regions (vs 0% of the recurrent variants, Fisher’s exact test, p=0.103, Figure 2a) and 21.1% of the unique protein-coding variants induced a non-synonymous change (vs 16.7% of the recurrent variants, Fisher’s exact test, p=1, Figure 2a). On a per-gene base, *MT-ND6* had lower mutation rates per base in the unique than in the recurrent variants (Fisher’s exact test, p=0.038, Figure 2b). The unique variants had lower heteroplasmic loads than the recurrent ones, with 80.0% of the unique variants having loads below 5% (vs 16.7% of recurrent variants with loads <5%, Chi-square test, p<0.0001, Figure 2c). In the unique variants, we found that the rRNA and tRNA variants were exclusively found at loads <5% (Figure 2h), and that the incidence of non-synonymous protein-coding variants with loads >10% was slightly higher than that of synonymous variants (Fisher’s exact test, p=0.043, Figure 2h).

Overall, these results show that half of oocytes differ from their siblings within a cohort due to the presence of variants unique to them. These unique variants differ from the recurrent variants in their location and load, being more frequently located in the rRNA and tRNA genes, and more often resulting in a non-synonymous change. Conversely, their pathogenic potential may be limited by their low heteroplasmic load, which in most cases was below 5%.

### The differences between unique and recurrent variants in oocytes are maintained during preimplantation development

Figure 3a shows an example of the variants found in each of the single blastomeres of two sibling embryos. The variants were categorized as recurrent across sibling embryos, recurrent across blastomeres (but not across the siblings) and unique to each blastomere. We reasoned that, further in development, the variants that are recurrent across oocytes of one donor become variants recurring across sibling embryos. Six variants of this type were found in nine embryos from the 4 sets of sibling day-3 embryos, with a similar distribution and heteroplasmic load as the recurrent variants we found in the oocytes. Half were located in the non-coding region and the other half in the protein-coding region, 16.7% being non-synonymous changes, the highest mutation rate per base being in the hypervariable region and 66.7% of variants having a heteroplasmic load over 20% (Figure 3b, Figure 3d and Figure 3f).

**Figure 3.**
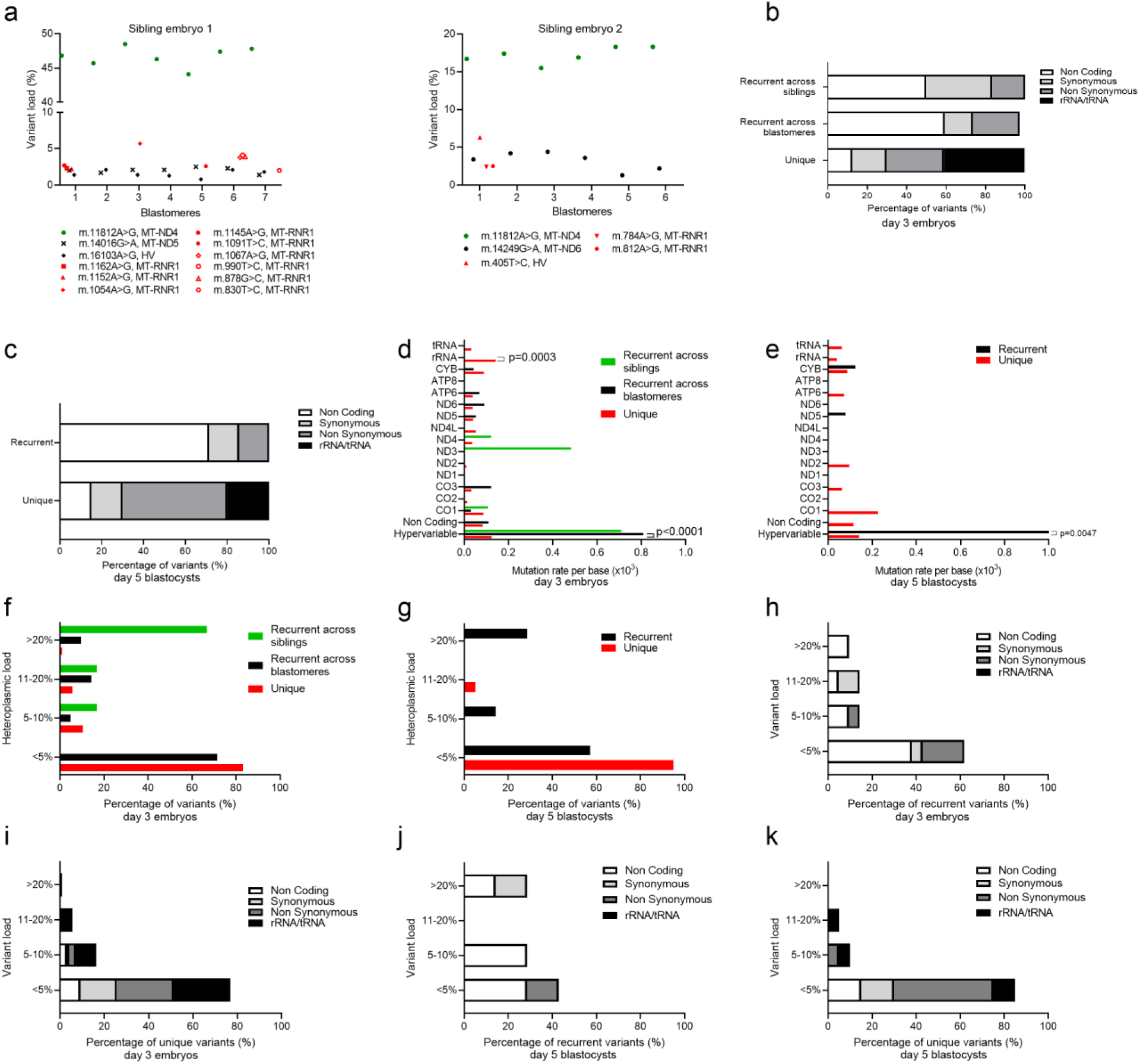
The differences in unique and recurrent variants are further maintained during preimplantation development. **a**. Example of two sibling embryos carrying recurrent variants across siblings (green dots), recurrent variants across blastomeres of the same embryo but not recurring across siblings (black dots) and unique variants that are unique to one blastomere (red symbols). **b**. Distribution of recurrent across siblings, recurrent across blastomeres and unique variants in day-3 embryos based on their location or type. **c**. Distribution of recurrent and unique variants in day-5 blastocysts based on their location or type. **d**. Mutation rate per base for the recurrent across siblings, recurrent across blastomeres and unique variants in the day-3 embryos. Variants in the hypervariable regions were more likely to be recurrent across blastomeres (Fisher’s exact test, p<0.0001) and variants in the rRNA regions were more likely to be unique (Fisher’s exact test, p=0.0003). **e**. Mutation rate per base for the recurrent and unique variants in the day-5 blastocysts. Variants in the hypervariable regions were more likely to be recurrent (Fisher’s exact test, p=0.0047). **f**. Recurrent across siblings, recurrent across blastomeres and unique variants in the day-3 embryos categorized for their variant load. **g**. Recurrent and unique variants in the day-5 blastocysts categorized for their variant load. **h**. Variants recurrent across blastomeres in the day-3 embryos categorized for their load and distribution in the mtDNA. **i**. Unique variants in the day-3 embryos categorized for their load and distribution in the mtDNA. **j**. Recurrent variants in the day-5 blastocysts categorized for their load and distribution in the mtDNA. **i**. Unique variants in the day-5 blastocysts categorized for their load and distribution in the mtDNA.

The second type of recurrent variant, which appears in multiple cells of the same day-3 embryo, but not between embryos of the same cohort, is found in 60% of embryos. These variants were located in 61.9% of cases in the non-coding region, 38.1% were in protein-coding sequences, 23.8% inducing non-synonymous changes, and none were found in the rRNA/tRNA regions (Figure 3b). The mutation rate per base was highest for the hypervariable region (Figure 3d). This distribution was maintained in the day-5 blastocysts, with 71.4% of variants in the non-coding region, 28.6% in the protein-coding regions, and none in the rRNA/tRNA, and with the highest mutation rate per base in the hypervariable region (Figure 3c and Figure 3e). On day 3, 9.5% of recurrent variants had loads >20% and 71.4% <5%, of which most were present in the non-coding regions (Figure 3f and Figure 3h). Noticeably, the recurrent protein-coding variants inducing non-synonymous changes were exclusively seen at loads <10%, while synonymous variants often showed loads >10% (Figure 3h). This distribution changed slightly on day 5, with 28.6% of recurrent variants having loads >20% and 57.1% <5%, the majority of which located in the non-coding regions (Figure 3g and Figure 3j). Remarkably, in both stages of development, these recurrent variants showed heteroplasmic loads that differed in average 3.5% across cells or biopsies, and maximally in 13.4%. This consistency is in line with previous reports on inherited pathogenic mtDNA variants detected during preimplantation genetic testing^39^. This shows that during the very early stages of development, these recurrent variants are homogeneously distributed to the daughter cells, suggesting that they were evenly distributed in the cytoplasm of the original oocyte. These variants restricted to one embryo differ from the ones found recurrently across embryos of the same cohort in their heteroplasmic load (71.4% of the recurrent variants across blastomeres have loads <5% vs 66.7% of the recurrent variants across siblings have loads >20%) but not in their location.

With regards to the unique variants, the difference in location was more prominent in the day-3 embryos, where 41.0% of the unique variants were located in the rRNA/tRNA regions as compared to none of the recurrent variants (Fisher’s exact test, p<0.0001, Figure 3b). This was also seen when looking at the mutation rate per base, where the mutation rate of the hypervariable region was significantly higher in the recurrent variants than in the unique (Fisher’s exact test, p<0.0001) and the rRNA regions showed higher rates in the unique variants (Fisher’s exact test, p=0.0003, Figure 3d). On day 3, 83.1% of the unique variants and 71.4% of the recurrent variants had a heteroplasmic load <5% (Chi-square test, p<0.0001, Figure 3f). Most of the variants with a load <5% were protein-coding variants with a similar distribution between synonymous and non-synonymous variants (Figure 3i). On day 5 of development, the variants in the hypervariable region were more likely to be recurrent (Fisher’s exact test, p=0.0047, Figure 3e) while the unique rRNA/tRNA variants represented only 20.0% of the variants (Fisher’s exact test, p=0.55, Figure 3c). The heteroplasmic loads were still different though not significant due to the limited sample size, with 85.0% of the unique variants and 57.1% of the recurrent variants having loads <5% (Fisher’s exact test, p=0.29, Figure 3g), the majority of which located in the protein-coding regions inducing non-synonymous changes (Figure 3k).

### The cells carrying unique mtDNA variants in embryos likely give rise to stable lineages in adult individuals

In 38.7% of the bulk adult tissues, we found variants that recurred between at least two samples of the same individual. These variants resembled the recurrent variants found in the oocytes and across sibling embryos, with 45.5% located in the non-coding regions, 48.5% in the protein-coding sequences and only a small proportion in the rRNA/tRNA coding regions (6.1%) (Figure 4a). The mutation rate per base was highest in the hypervariable region (Figure 4b), and most frequently had loads >20% (Figure 4c). Further, there were no differences in the type of variants in function of their variant load (Figure 4d). Also, 16.7% of the samples carried unique variants. Of these, 66.7% were found in the non-coding regions while no variants were located in the rRNA/tRNA regions (Figure 4a), and the mutation rate per base was as expected highest in the hypervariable region (Figure 4b). Of the unique variants, 85.2% had loads <5%, which was significantly different compared to 24.2% of the recurrent variants (Fisher’s exact test, p<0.0001, Figure 4c), of which 65.2% were located in the non-coding regions (Figure 4e). In sum, the only difference between the recurrent and unique variants in the bulk DNA samples was their heteroplasmic load.

**Figure 4.**
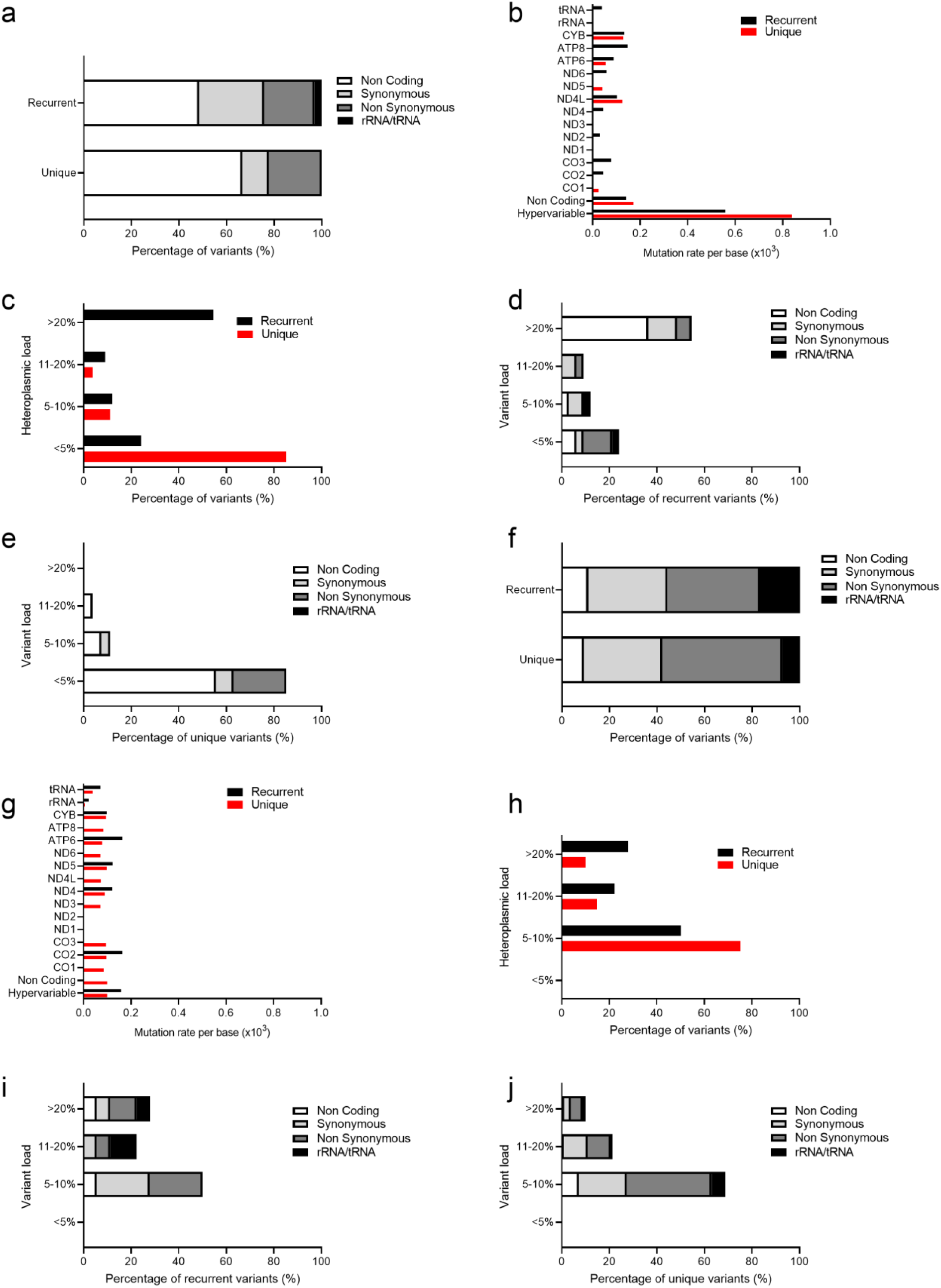
The cells carrying unique mtDNA variants in embryos give rise to stable lineages in adult individuals. **a**. Distribution of recurrent and unique variants in adult bulk tissues based on their location or type. **b**. Mutation rate per base for the recurrent and unique variants in adult bulk tissues. **c**. Recurrent and unique variants in adult bulk tissues categorized for their variant load. **d**. Recurrent variants in adult bulk tissues categorized for their load and distribution in the mtDNA. **e**. Unique variants in adult bulk tissues categorized for their load and distribution in the mtDNA. **f**. Distribution of recurrent and unique variants in adult single cells based on their location or type. **g**. Mutation rate per base for the recurrent and unique variants in adult single cells. **h**. Recurrent and unique variants in adult single cells categorized for their variant load. **i**. Recurrent variants in adult single cells categorized for their load and distribution in the mtDNA. **j**. Unique variants in adult single cells categorized for their load and distribution in the mtDNA.

Next, we studied single cells of two tissues of three individuals. In this part, we set the threshold for variant calling at >5% to ensure a more conservative calling. The main reason for this is the higher number of PCR cycles required to amplify the mtDNA of these cells, which could result in an increase in false positives. Of the adult single cells, 47.1% carried recurrent mtDNA variants (found in at least two different cells from one individual, irrespective of the tissue of origin), while unique variants appeared in 85.3% of cells. In total, we identified 19 recurrent variants and 642 unique variants in the 68 single cells. Furthermore, not all recurrent variants in the single cells were identified in the bulk samples of the same tissues, as the average of the loads of all cells could drop below the sequencing detection limit.

The location of the recurrent and unique variants in the adult single cells was very similar, with most variants located in coding and tRNA/rRNA loci, and this in contrast with the variants identified in the bulk tissues. Protein-coding variants were more frequent at the single-cell level than in the bulk tissues (recurrent bulk 48.5% vs single-cell 68.4%, Chi-square, p<0.0001, unique bulk 33.3%. vs single-cell 83.3%, Chi-square, p<0.0001) reminiscent of the unique variants in the day-3 and day-5 blastocysts, suggesting that the cells carrying unique variants in the preimplantation embryos can give rise to stable lineages in terms of mtDNA variants in the adult individual. In the same line, there were no differences between the recurrent and unique single-cell variants in mutation rate per base across the different loci (Figure 4g) but these were significantly different to their counterparts in bulk tissues, where the hypervariable region had the highest mutation rate per base. As in the bulk tissues, recurrent variants tended to have higher variant loads than the unique variants, and no differences were found in location according to heteroplasmic load (Figure 4h, Figure 4i, Figure 4j). Finally, we controlled the recurrent and unique single-cell variants for the type of base pair change they induce, and found no differences in the incidence of transitions, transversions or insertions and deletions (Figure S1).

## DISCUSSION

In this study we deep-sequenced the mtDNA of a large cohort of human oocytes, blastomeres of day-3 embryos, small groups of cells of day-5 blastocysts, and single cells as well as bulk DNA of different somatic adult tissues. We identified three different types of mtDNA variants based on their recurrence or uniqueness across siblings, single cells or samples from the same individuals. These types of variants are differently distributed throughout the regions of the mtDNA and show differences in variant load.

In early development, the similarities between the low-load unique variants in the oocytes and the unique variants in the day-3 and day-5 blastocysts on one hand, and between the recurrent variants in the oocytes and recurrent across sibling embryos on the other hand leads us to propose the following model of segregation for each type of variant (illustrated in Figure 5). The recurrent variants across oocytes are the equivalent of the recurrent variants across sibling embryos (Figure 5a), while the variants that are recurrent restricted to one embryo are the equivalent of the unique variants in oocytes that are found at higher heteroplasmic loads. We propose that both these variants are homogeneously distributed in mitochondria of the ooplasm, and therefore homogeneously segregated in the blastomeres throughout the cleavages (Figure 5b).

**Figure 5.**
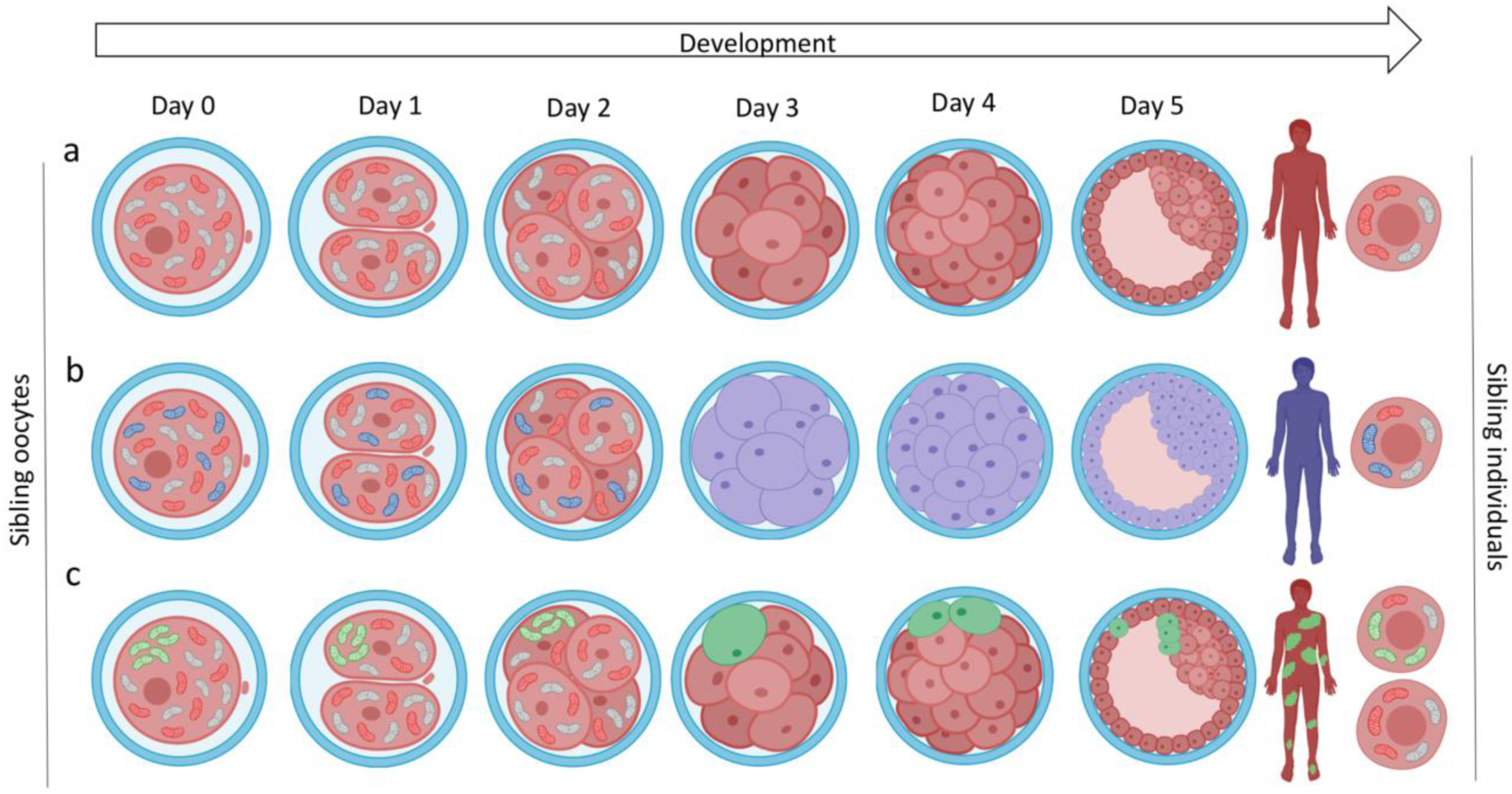
Proposed segregation model of recurrent and unique variants in the preimplantation development. The figure shows the development of three oocytes of the same donor, resulting in three sibling individuals. Mitochondria in red (a-c) are carrying variants that are recurrent across sibling oocytes and individuals. The mitochondria in blue (b) carry unique variants at higher loads in the oocyte and are homogenously distributed together with the red mitochondria in the ooplasm. They remain in a similar distribution during development and become the variants found as unique in one individual, but present in all their tissues. The mitochondria in green (c) carry unique variants at lower loads and cluster in the ooplasm. These mitochondria will remain in close proximity to each other during the cleavage stages and will be present in only one blastomere. This cell will then give rise to a lineage of cells -that in the adult individual-manifests as a subpopulation of rare cells carrying this same variant, potentially across different germ layers. This figure was created with BioRender.

Finally, we hypothesize that variants that are unique to a single blastomere would originate from the low-load unique variants in the oocytes (Figure 5c). This can be explained if these variants are found in a low number of mitochondria that cluster together in the ooplasm. It is likely that these low-load unique variants in the oocyte arise postnatally during folliculogenesis due to replication of a subpopulation of mtDNA molecules, as demonstrated in a mouse model, while the high-load unique variants were already present in the primordial germ cells after the first bottleneck^40^. This would explain why the low-load unique variants in the oocytes (and unique across blastomeres) are more frequently non-synonymous and in rRNA/tRNA loci than the high-load unique variants in oocytes (and recurrent across blastomeres), since the latter were subjected to the selecting effect of the germline bottleneck^11^.

During the first embryonic cleavages, this co-localization would result in an asymmetric distribution of mitochondria containing the variant, leading to an embryo that already presents mosaicism as early as day 3 of development. This same pattern would continue through to the blastocyst stage, where sister cells tend to remain in proximity of each other. These lineages persist throughout development, and in the adult individual as supported by the study of Lee et al. in 2012^41^.

It is plausible that this type of segregation of mtDNA variants is a common event during the cleavage-stages of mammalian development. Work on mice and Rhesus monkeys have shown that this type of segregation indeed occurs in the preimplantation period of these two models^41,42^. Also, work on human embryos has shown that asymmetrical mitochondrial distribution can result in a proportion of blastomeres with a reduced mitochondrial pool^43^, which could contribute to the fixing of mtDNA variants in the clonally derived blastomeres.

Despite having identified the unique variants throughout development, it is noteworthy that there is a decline in unique rRNA and tRNA variants from day 3 to day 5 of development. This suggests the existence of selection mechanisms during preimplantation development that filter out pathogenic variants. This selection has been observed later in the mouse development, where deleterious variants present in the oocyte and in the embryo were found to be selected against by unknown mechanisms occurring postimplantation and postnatally^44^.

Next to possible selection mechanisms, the somatic bottleneck can cause a diverse population of cells harboring different mtDNA heteroplasmic variants and at different loads to exist. We saw that the variant load in adult single cells varies widely amongst cells of the same tissue. The number of cells harboring variants at higher loads could exceed a certain tissue threshold and could increase the risk of pathologies^45^. This was already described in muscle fibers^46^ and the heart^47^.

Furthermore, our study confirmed the presence of tissue-specific variants. These variants were located in the non-coding regions and were in close proximity to regions that regulate the mtDNA replication. This is in line with the suggestion of Samuels et al.^22^ that these variants could have a beneficial effect on the mtDNA replication in the given tissue. For instance, variant m.72T>C was already described by two other research groups in liver and kidney tissue^21,22^.

In conclusion, our work is the first to comprehensively describe mtDNA mosaicism in early human development and identifies a subgroup of low-load variants that may give rise to stable lineages of genetically diverse cells in the adult. We propose that these lineages appear due to asymmetrical distribution of mitochondria carrying mtDNA variants in the oocyte, which possibly appeared during folliculogenesis. Finally, future research will give us more insight on the mechanisms behind the asymmetrical distribution of variants in the oocyte and on the potential implications of this type of mosaicism in health and disease.

## SUPPLEMENTAL DATA

**Table S1.** Overview of sample characteristics included in this study.

|  | Sample size |
| --- | --- |
| **Oocytes** | 254 |
| Donors |  |
| Oocyte donors | 44.5% (113) |
| ART Patients | 55.5% (141) |
| Maturation stage |  |
| M1 | 20.1% (51) |
| GV | 37.4% (95) |
| M2 | 22.4% (57) |
| Unknown | 20.1% (51) |
| **Embryos** |  |
| **Day-3 embryos** | 25 |
| Donor couples | 9 |
| Blastomeres | 158 |
| Average number of blastomeres per embryo | 6.3 |
| Sibling embryos/pairs | 22 of 6 couples |
| **Day-5 blastocysts** | 7 |
| Donor couples | 6 |
| Inner Cell Mass | 7 |
| Trophectoderm | 10 |
| **Adult bulk tissues** |  |
| Donors | 60 |
| Age | 18-55 |
| Buccal | 59 |
| Blood | 57 |
| Urine | 26 |
| **Adult single cells** |  |
| Donors | 3 |
| Age donor 1 | 26 |
| Age donor 2 | 27 |
| Age donor 3 | 55 |
| Buccal | 37 |
| Urine | 31 |

**Table S2.**
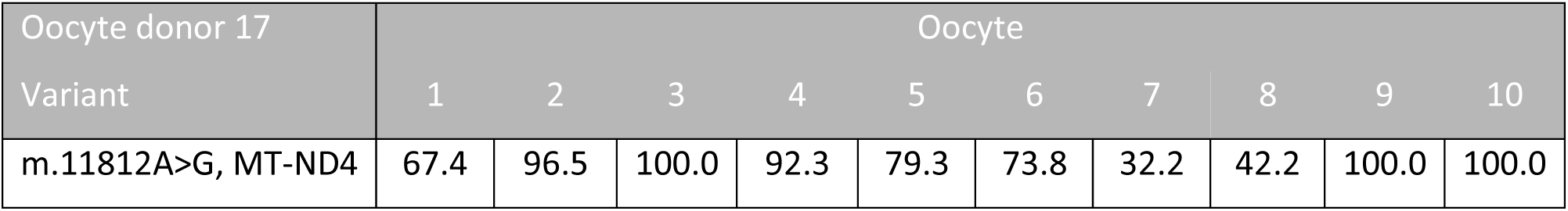
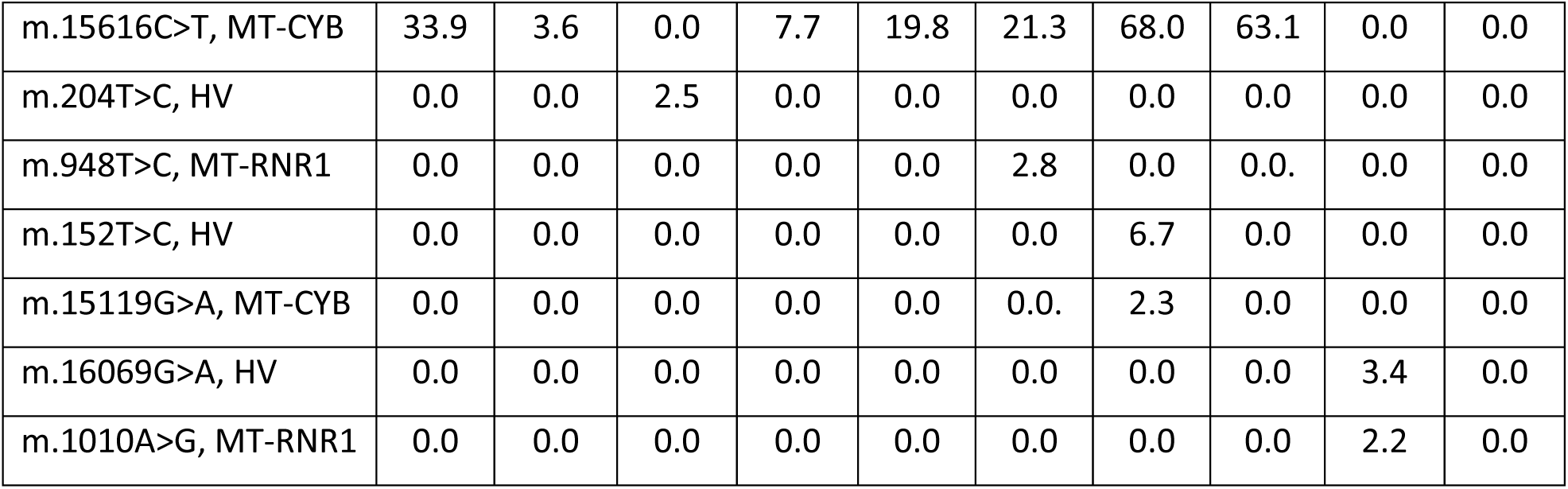
Example of the variant composition in multiple oocytes from one donor.

**Table S3.**
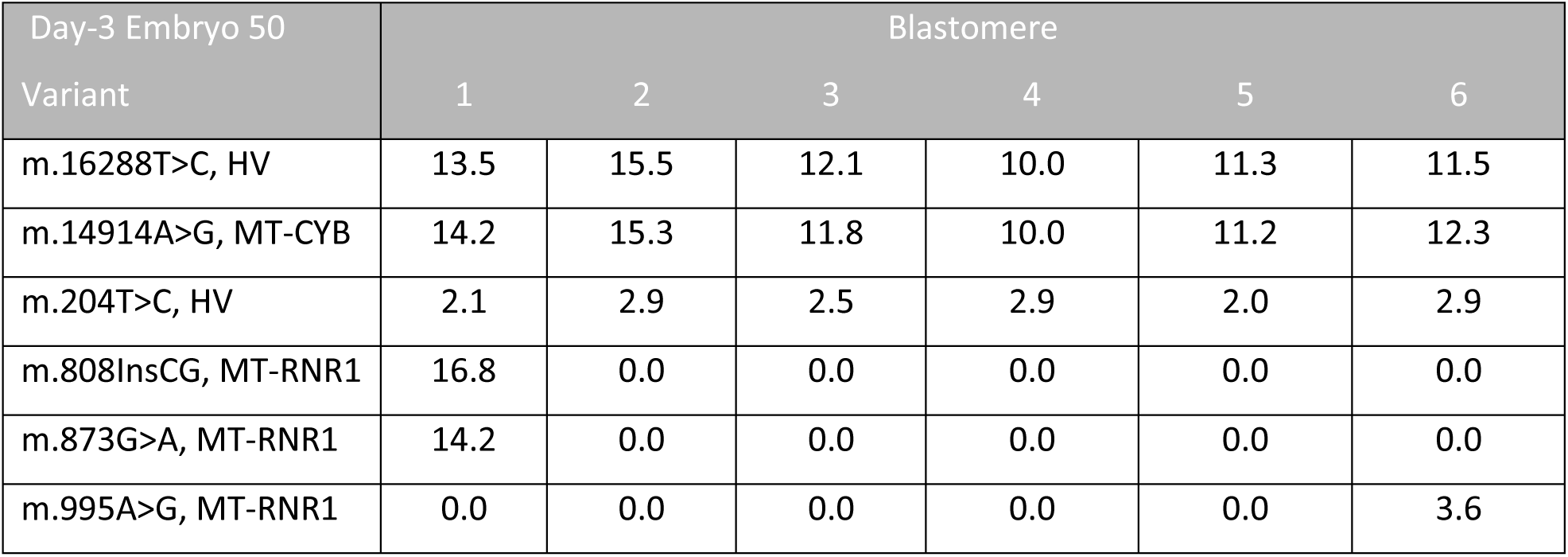
Example of the variant composition in multiple blastomeres from one day-3 embryo.

**Table S4.** Example of the variant composition in multiple biopsies from one blastocyst.

| Day-5 Embryo 3 |  |  |  |
| --- | --- | --- | --- |
| Variant | Inner Cell Mass | Trophectoderm 1 | Trophectoderm 2 |
| m.12831C>T, MT-ND5 | 23.1 | 27.8 | 22.2 |
| m.1097G>A, MT-RNR1 | 12.0 | 0.0 | 0.0 |
| m.6372T>C, MT-CO1 | 4.9 | 0.0 | 0.0 |
| m.6123A>G, MT-CO1 | 2.4 | 0.0 | 0.0 |
| m.6080A>G, MT-CO1 | 2.4 | 0.0 | 0.0 |
| m.5606C>T, MT-TA | 2.0 | 1.10 | 0.0 |
| m.9672C>T, MT-CO3 | 0.0 | 2.7 | 0.0 |
| m.8810C>T, MT-ATP6 | 0.0 | 2.1 | 0.0 |
| m.5890C>T, MT-TY | 0.0 | 0.0 | 5.2 |

**Table S5.** Example of the variant composition in multiple tissues from one individual.

| Adult Bulk Tissue |  |  |  |
| --- | --- | --- | --- |
| Oocyte donor 10 |  |  |  |
| Variant | Buccal | Blood | Urine |
| m.203G>A, HV | 37.4 | 5.3 | 28.43 |
| m.215A>G, HV | 3.2 | 0.0 | 0 |
| m.9525G>A, MT-CO3 | 1.5 | 4.0 | 2.16 |

**Figure S1.**
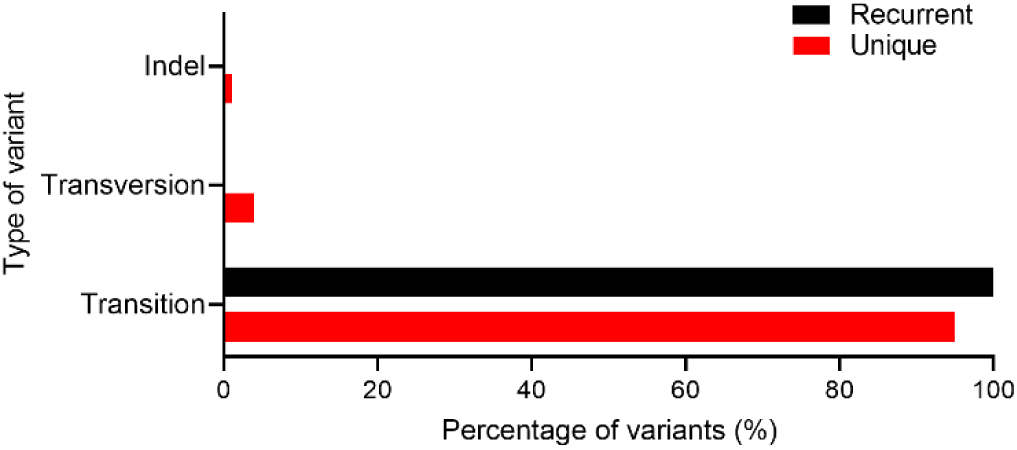
Number of transitions, transversions and insertions and deletions (indel), categorized for recurrent and unique, found in adult single cells. No differences were found in the different cohorts.

## WEB RESOURCES

mtDNA server https://mitoverse.i-med.ac.at/index.html#!run/mtdna-server%40v2.0.0

MitoWheel http://www.mitowheel.org/mitowheel.html

MutPred2 http://mutpred.mutdb.org/

BioRender https://biorender.com/

## ACKNOWLEDGEMENTS

This work was supported by the Scientific Fund Willy Gepts of the UZ Brussel (Wetenschappelijk Fonds Willy Gepts), the Methusalem Grant of Karen Sermon of the Vrije Universiteit Brussel and the Research Foundation Flanders (Fonds voor Wetenschappelijk Onderzoek Vlaanderen, FWO 1518418N). M.R. and E.C.D.D. are doctoral fellows at the FWO.

## DATA AND CODE AVAILABILITY

The dataset supporting the current study has not been deposited in a public repository because the participants who donated their genetic material did not agree to share their personal sequencing information when they signed informed consent. However, researchers can access the data from the corresponding author on request for further legal use. The data will be formatted in an anonymized Microsoft Excel file where only the type of heteroplasmic variant, the variant load, the tissue of origin and if the variant was recurrent across samples will be available.

